# Activity-dependent stabilization of nascent dendritic spines requires non-enzymatic CaMKIIα function

**DOI:** 10.1101/2022.07.18.500536

**Authors:** Nicole Claiborne, Karen Zito

**Affiliations:** Center for Neuroscience, University of California, Davis, CA 95618

**Keywords:** dendritic spine, CaMKII, two-photon imaging, glutamate uncaging

## Abstract

The outgrowth and stabilization of nascent dendritic spines are crucial processes underlying learning and memory. Most new spines retract shortly after growth; only a small subset is stabilized and integrated into the new circuit connections that support learning. New spine stabilization has been shown to rely upon activity-dependent molecular mechanisms that also contribute to long-term potentiation (LTP) of synaptic strength. Indeed, disruption of the activity-dependent targeting of the kinase CaMKIIα to the GluN2B subunit of the NMDA-type glutamate receptor disrupts both LTP and activity-dependent stabilization of new spines. Yet it is not known which of CaMKIIα’s many enzymatic and structural functions are important for new spine stabilization. Here, we used two-photon imaging and photolysis of caged glutamate to monitor the activity-dependent stabilization of new dendritic spines on hippocampal CA1 neurons from mice of both sexes in conditions where CaMKIIα functional and structural interactions were altered. Surprisingly, we found that inhibiting CaMKIIα kinase activity either genetically or pharmacologically did not impair activity-dependent new spine stabilization. In contrast, shRNA knock-down of CaMKIIα abolished activity-dependent new spine stabilization, which was rescued by co-expressing shRNA-resistant CaMKIIα. Notably, overexpression of phospho-mimetic CaMKIIα-T286D, which exhibits activity-independent targeting to GluN2B, enhanced basal new spine survivorship in the absence of additional glutamatergic stimulation, even when kinase activity was disrupted. Together, our results suggest that nascent dendritic spine stabilization requires structural and scaffolding interactions mediated by CaMKIIα that are independent of its enzymatic activities.

**SIGNIFICANCE STATEMENT:** The stabilization of nascent dendritic spines is thought to support lasting memory of learned experiences. Here, we show that scaffolding and structural interactions, but not the enzymatic activities, of the kinase CaMKIIα are required for activity-dependent new spine stabilization. This study furthers our understanding of the cellular and molecular processes that facilitate learning and memory in the mammalian brain. Understanding the cellular and molecular mechanisms of learning and memory is crucial for our ability to develop therapeutics for memory impairments associated with neurological and neurodegenerative disorders.

## INTRODUCTION

The dynamic modification of neuronal circuitry underlies learning and memory and is crucial for adaptation and survival. Dendritic spines are the sites of most excitatory synaptic connections in the mammalian cerebral cortex, and the morphological and functional changes that occur at dendritic spines contribute to the neural circuit modifications that support behavior (Kasai et al., 2021). Notably, the stabilization of newly formed spines in the cortex is tightly linked to lasting memory of learned experiences (Hayashi-Takagi et al., 2015; Roberts et al., 2010; Xu et al., 2009; Yang et al., 2009). Interestingly, most new spines are transient (Berry and Nedivi, 2017; Holtmaat et al., 2005) suggesting that stabilization is precisely regulated to favor only a subset of new spines sufficient to support memory. Thus, defining the mechanisms that determine which new spines are stabilized will strengthen our understanding of learning and memory.

Previous studies have shown that synaptic activity enhances the stability of new dendritic spines in the hippocampus, and that the enhancement of new dendritic spine stability appears to be specific to patterns of synaptic activity that result in the coordinated long-term enhancement of synaptic strength and spine volume (Matsuzaki et al., 2004) known as long-term potentiation (LTP) (De Roo et al., 2008a; Hill and Zito, 2013). NMDA-type glutamate receptor (NMDAR) activation is required for LTP-induced nascent spine stabilization, and disruption of the interaction between the Ca^2+^/calmodulin-activated kinase CaMKIIα and the GluN2B subunit of the NMDAR prevents activity-dependent new spine stabilization (Hill and Zito, 2013). Notably, CaMKIIα-GluN2B binding facilitates a number of CaMKIIα enzymatic and structural functions that promote LTP induction and maintenance, including binding to densin-180 and α-actinin, activation of signaling molecules, and phosphorylation of AMPA-type glutamate receptors (Bayer and Schulman, 2019; Sanhueza and Lisman, 2013). Whether these enzymatic and structural activities of CaMKIIα and the downstream cascades they initiate are required for activity-dependent new spine stabilization is not yet known.

Here, we used time-lapse imaging and two-photon glutamate uncaging along with genetic and pharmacological manipulations to elucidate the role of CaMKIIα in activity-dependent new spine stabilization. We first demonstrated that CaMKIIα is present and enriched at mature levels in new spines shortly after outgrowth on CA1 neurons in hippocampal slice cultures, supporting that CaMKIIα could play an important role in nascent spine stabilization. Surprisingly, we found that high-frequency glutamate uncaging (HFU) enhanced new spine survivorship even when CaMKIIα kinase activity was genetically and pharmacologically inhibited. In contrast, shRNA-mediated knock-down of CaMKIIα blocked activity-dependent new spine stabilization, indicating that CaMKIIα is indeed required for new spine stabilization. Finally, we show that autonomous, constitutively-active CaMKIIα-T286D enhanced new spine stability without further glutamatergic stimulation or kinase activity. Together, our results support a model whereby strong glutamatergic transmission at a subset of new spines facilitates new spine stabilization through structural and scaffolding functions of CaMKIIα.

## METHODS

### Preparation and transfection of organotypic slice cultures

Organotypic hippocampal slice cultures were prepared from postnatal day (P) 6-8 C57BL/6J wild-type mice of both sexes, as described (Opitz-Araya and Barria, 2011; Stoppini et al., 1991). Neurons were transfected 2-3 days prior to imaging using particle-mediated gene transfer, as described (Woods and Zito, 2008), except 6-8 µg of DsRed-Express (Clontech) and 6 μg of mEGFP-tagged constructs or 5-10 μg of mEGFP were coated onto 6-7 mg of 1.6 μm gold beads. mEGFP-tagged constructs included: GFP-CaMKIIα, GFP-CaMKIIα-T286D, GFP-CaMKIIα-K42R/T286D (Pi et al., 2010), or GFP-CaMKIIα-K42R (Tullis et al., 2020). CaMKIIα knockdown used 25 µg CaMKIIα-shRNA and rescue also contained 6 µg shRNA-resistant mEGFP-CaMKIIα* (Lemieux et al., 2012).

### Two-photon imaging

Image stacks (512 × 512 pixels, 1 μm z-steps) of 4-6 secondary and tertiary, apical and basal dendritic segments from CA1 pyramidal neurons (6-10 DIV) were acquired on a custom two-photon microscope with a pulsed Ti::Sapphire laser (930 nm, 0.5-3 mW at the sample; Spectra Physics, Newport). Data acquisition was controlled by ScanImage (Pologruto et al., 2003) written in MATLAB (MathWorks). All images shown are maximum projections of 3D image stacks after applying a median filter (3 × 3). The first time point was acquired in slice culture medium at room temperature (RT) and the slice was maintained in the incubator (35°C) for 1 h between first and subsequent acquisitions. After 1 h, the slice was placed in a bath of recirculating, oxygenated artificial cerebrospinal fluid (ACSF) (in mM: 127 NaCl, 25 NaHCO_3_, 1.2 NaH_2_PO_4_, 2.5 KCl, 25 D-glucose, ∼310 mOsm, pH 7.2) with 2 mM Ca^2+^, 0-0.1 mM Mg^2+^, and 1 *μ*M tetrodotoxin at 31°C. 2.5-3.5 mM of 4-methoxy-7-nitroindolinyl-caged L-glutamate (MNI-glutamate) was added for uncaging experiments. Staurosporine (1 *μ*M) or an equivalent volume vehicle were added to the bath after new spine identification and 30 min prior to uncaging.

### Identification new spines and estimation of spine size

We defined new spines as any protrusion emanating from the dendrite that was present in the second and/or third images in the time-lapse series (60-90 min later) but not detectable in either the red or green channels in the first image. Persistent neighbor spines were defined as spines that were present in all images in the time-lapse series. Spines of ambiguous persistence or presence due to fluctuations in dendrite swelling, spine motility, or spine drift in the z-axis were excluded. Spine size was estimated from bleed-through-corrected and background-subtracted red (DsRed-Express) fluorescence intensity. Spine brightness measurements give an accurate estimate of relative spine size when compared with electron microscopy (Holtmaat et al., 2005).

### High frequency uncaging (HFU) stimulus

The HFU stimulus consisted of 60 pulses (720 nm, 8-10 mW at the sample) of 2 ms duration delivered at 2 Hz in the presence of 2.5-3.5 mM MNI-glutamate by parking the beam at a point ∼0.5 μm from the spine head away from the dendrite.

### Quantification of relative enrichment of GFP-tagged proteins

Relative enrichment of GFP-tagged proteins in dendritic spines was calculated using bleed-through-corrected and background-subtracted green (GFP) and red (DsRed-Express) fluorescence intensities from spines and dendrites, as described (Woods et al., 2011). Briefly, the ratio of green fluorescence intensity to red fluorescence intensity (G/R) was calculated for each new spine, size-matched neighboring persistent spines (6-10), and three representative regions on the dendritic shaft (excluding regions dendrite swelling and GFP-puncta, which were indicative of the presence of a z spine). To quantify spine fluorescence intensities, boxes were drawn around whole spines and spine necks using custom software written in MATLAB. Background subtraction was done by drawing a box next to a target spine that was equal on the axis perpendicular to the dendrite as the box drawn around the spine head and neck. The average intensity of that box was multiplied by the number of pixels in the target spine box and subtracted from the integrated intensity from the target spine box. Relative enrichment of spines was calculated by normalizing the G/R ratio of the target spine to the mean G/R ratio of three locations on the adjacent dendrite.

Several criteria were used to ensure that analyzed data were of high quality. Cells that exhibited lower green fluorescence intensity than the background ROI were excluded. Cells with extremely high levels of GFP-tagged protein expression such that synaptic enrichment was lost were excluded. Cells were also excluded if, after background and bleed-through subtraction: (1) the value of the mean green pixel intensity (G) from neighbor spines was less than 3.23 a.u., (2) the value of the mean neighbor spine G/R was less than 0.01, or (3) the ratio of the square of the mean persistent spine G/R to the absolute value of the mean dendrite G/R was less than 0.05. These criteria allowed unbiased exclusion of cells that returned negative pixel intensity values after background and bleed-through subtraction. Cells that exhibited significant photobleaching (a decline in average integrated fluorescence intensity in the dendrite greater than 20% compared to the first time point) in either the red or green channels were excluded.

### Statistical Analysis

Survivorship curves were compared using the log-rank task. To compare survivorship at individual time points, we used Fisher’s exact test. For comparisons of spine volumes at a given time point to baseline, 2-way ANOVAs with appropriate *post-hoc* tests for multiple comparisons were used. Between-group comparisons of spine baseline volume were performed using a two-tailed unpaired heteroscedastic Student’s *t* test unless otherwise noted. Error bars represent standard error of the mean (SEM).

## RESULTS

### GFP-CaMKIIα enrichment in new spines is comparable to that in size-matched neighboring spines

To understand the role of CaMKIIα in activity-induced new spine stabilization, we first needed to determine whether CaMKIIα is expressed in new spines and in what time frame. This experiment was an important first step, as we and others have reported that several members of the PSD-MAGUK family of postsynaptic scaffolding molecules are expressed at very low levels in new spines and can take up to 24 h to accumulate to mature levels (De Roo et al., 2008a; Lambert et al., 2017), indicating that the molecular composition of new spines and their persistent neighbors is distinct, particularly in the earliest stages after new spine outgrowth. We used time-lapse, 2-photon imaging to observe spontaneous new spine outgrowth on the dendrites of hippocampal CA1 neurons in slice culture biolistically-transfected with mEGFP-tagged CaMKIIα (GFP-CaMKIIα) and a DsRed-Express cell fill (**Fig. 1A**). We found no difference in the enrichment of GFP-CaMKIIα in new spines as compared to size-matched neighboring control spines (new: 1.5 ± 0.2; neighbor: 1.7 ± 0.1; *p = 0*.*14*; **Fig. 1B, C**). We conclude that CaMKIIα rapidly accumulates at new spines and therefore could play an important role in even the earliest molecular signaling events that support new spine stabilization.

**Figure 1.**
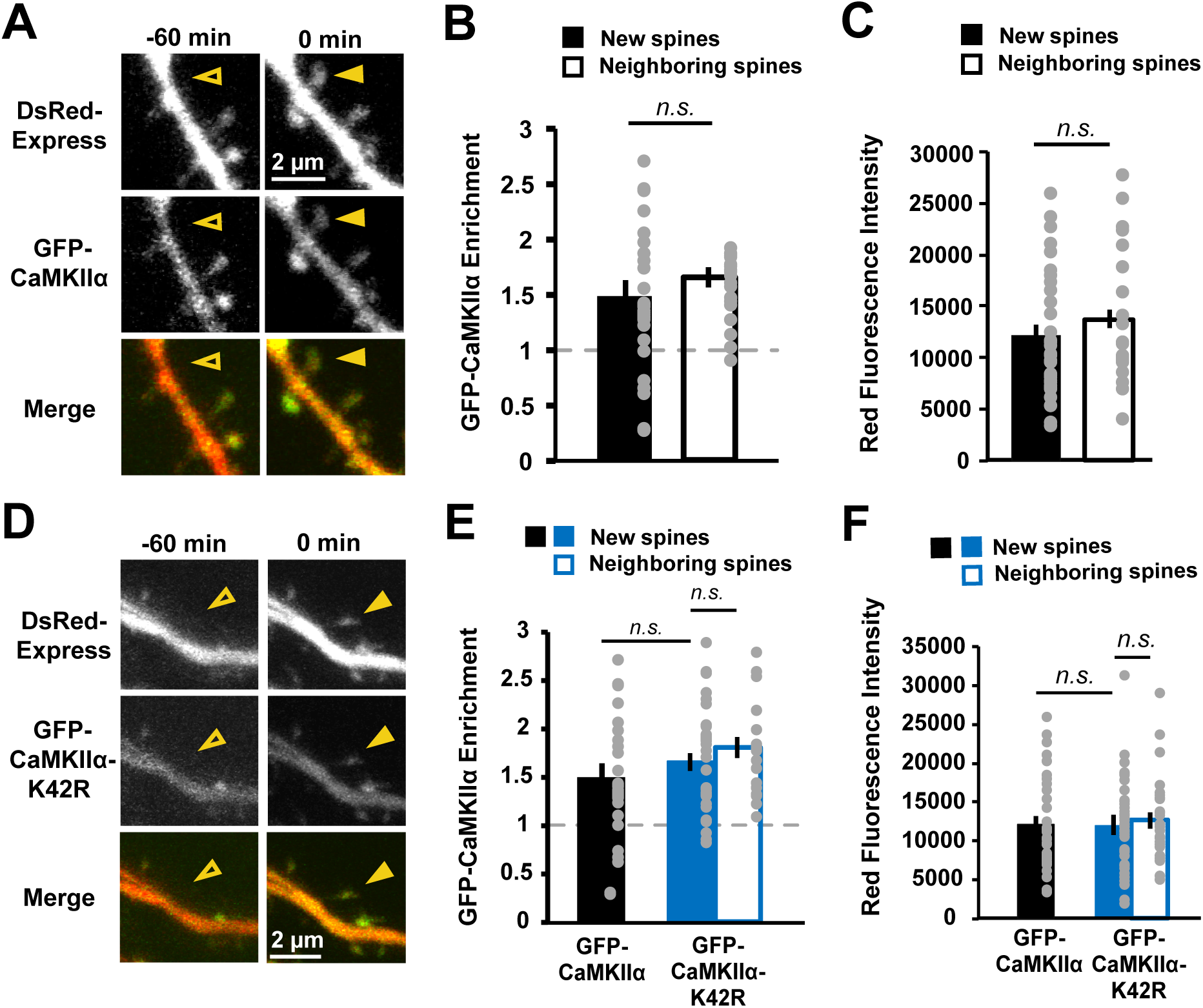
GFP-CaMKIIα enrichment in new spines is comparable to that in size-matched neighboring spines. ***(A)*** Images of dendrites from hippocampal CA1 neurons in slice culture (DIV 7-9) expressing GFP-CaMKIIα (green) and DsRed-Express (red) before (open arrowhead) and after (filled arrowhead) spontaneous new spine outgrowth. ***(B)*** Enrichment (spine: dendrite ratio) of GFP-CaMKIIα in new spines (n = 33 spines/16 cells) was comparable to that in size-matched neighboring spines (n = 21 spines/16 cells). ***(C)*** Neighboring spines used for enrichment calculations in ***B*** were size-matched to new spines (*p = 0*.*62*). ***(D)*** Images of dendrites from CA1 neurons expressing GFP-CaMKIIα-K42R (green) and DsRed-Express (red) before (open arrowhead) and after (filled arrowhead) spontaneous new spine outgrowth. ***(E)*** Enrichment of GFP-CaMKIIα-K42R in new spines (n = 39 spines/ 14 cells) was comparable to that in size-matched mature neighboring spines (n = 39 spines/14 cells). Importantly, no difference in relative enrichment was found between new (filled bars) or size-matched neighboring spines (open bars) in the WT (black) and K42R (blue) conditions (new: *p = 0*.*37*; neighbors: *p=0*.*99*). Data for CaMKIIα-WT new spine enrichment is from ***B. (F)*** Neighboring spines used for enrichment calculations in ***E*** were size-matched to new spines (*p = 0*.*99*). No difference in new spine size between WT (black) and K42R (blue; *p=0*.*99*). Data for WT new spine size is from ***C***. Two-way ANOVA with Bonferroni multiple comparisons test. *p <0.05, **p < 0.01, ***p < 0.001.

### Genetic and pharmacological inhibition of CaMKIIα kinase activity does not impair activity-dependent new spine stabilization

To investigate the role of CaMKIIα in activity-dependent new spine stabilization, we tested whether interfering with CaMKIIα kinase activity would disrupt the robust activity-dependent stabilization of new spines induced by high frequency uncaging (HFU) of MNI-caged glutamate (MNI-glutamate) at individual new spines (Hill and Zito, 2013). We first chose to use a genetic approach by overexpressing GFP-CaMKIIα containing the K42R point mutation that inhibits CaMKIIα kinase activity (Pi et al., 2010; Tullis et al., 2020; Yamagata et al., 2009). This CaMKIIα-K42R mutant has been shown to act in a dominant-negative manner (Pi et al., 2010; Rossetti et al., 2017). Importantly, enrichment of GFP-CaMKIIα-K42R in new spines was comparable to that in size-matched mature neighboring spines (new: 1.7 ± 0.1; neighbor: 1.8 ± 0.1; *p = 0*.*39*) and basal spine enrichment levels of GFP-CaMKIIα-K42R in new spines were comparable to those of GFP-CaMKIIα (*p = 0*.*37*; **Fig. 1D-F**).

We proceeded to test whether expression of GFP-CaMKIIα-K42R would disrupt stabilization of nascent dendritic spines. We used time-lapse imaging of dendrites on neurons expressing dsRed-Express and either GFP-CaMKIIα or GFP-CaMKIIα-K42R to identify multiple new spines that spontaneously grew on each cell. One new spine per cell was exposed to HFU stimulation (**Fig. 2A**). Survivorship of stimulated and unstimulated new spines on the same cell was monitored through time-lapse imaging. Consistent with our observations for cells transfected with GFP alone (Hill and Zito, 2013), our HFU protocol significantly enhanced stimulated new spine survivorship compared to unstimulated new spines on cells expressing GFP-CaMKIIα (**Fig. 2B-D**; stim: 94%, unstim: 62%; *p = 0*.*03*). Surprisingly, we found that HFU also robustly enhanced new spine stabilization on cells expressing GFP-CaMKIIα-K42R (**Fig. 2B-D**; stim: 100%, unstim: 68%; *p = 0*.*02*), suggesting that CaMKIIα kinase activity is not necessary for activity-induced new spine stabilization. Indeed, the rate of stimulated and unstimulated new spine survivorship were not different between the GFP-CaMKIIα or GFP-CaMKIIα-K42R conditions (stim GFP-CaMKIIα vs GFP-CaMKIIα-K42R: *p = 0*.*99*; unstim GFP-CaMKIIα or GFP-CaMKIIα-K42R: *p = 0*.*79*). Importantly, we confirmed that GFP-CaMKIIα-K42R was acting as a dominant negative, as GFP-CaMKIIα-K42R-transfected neurons exhibited impaired HFU-induced long-term growth of mature spines (K42R: 138 ± 16%; *p = 0*.*11*), which is intact in neurons expressing GFP-CaMKIIα (WT: 188 ± 23%; *p = 0*.*01*; **Fig. 2E, F**).

**Figure 2.**
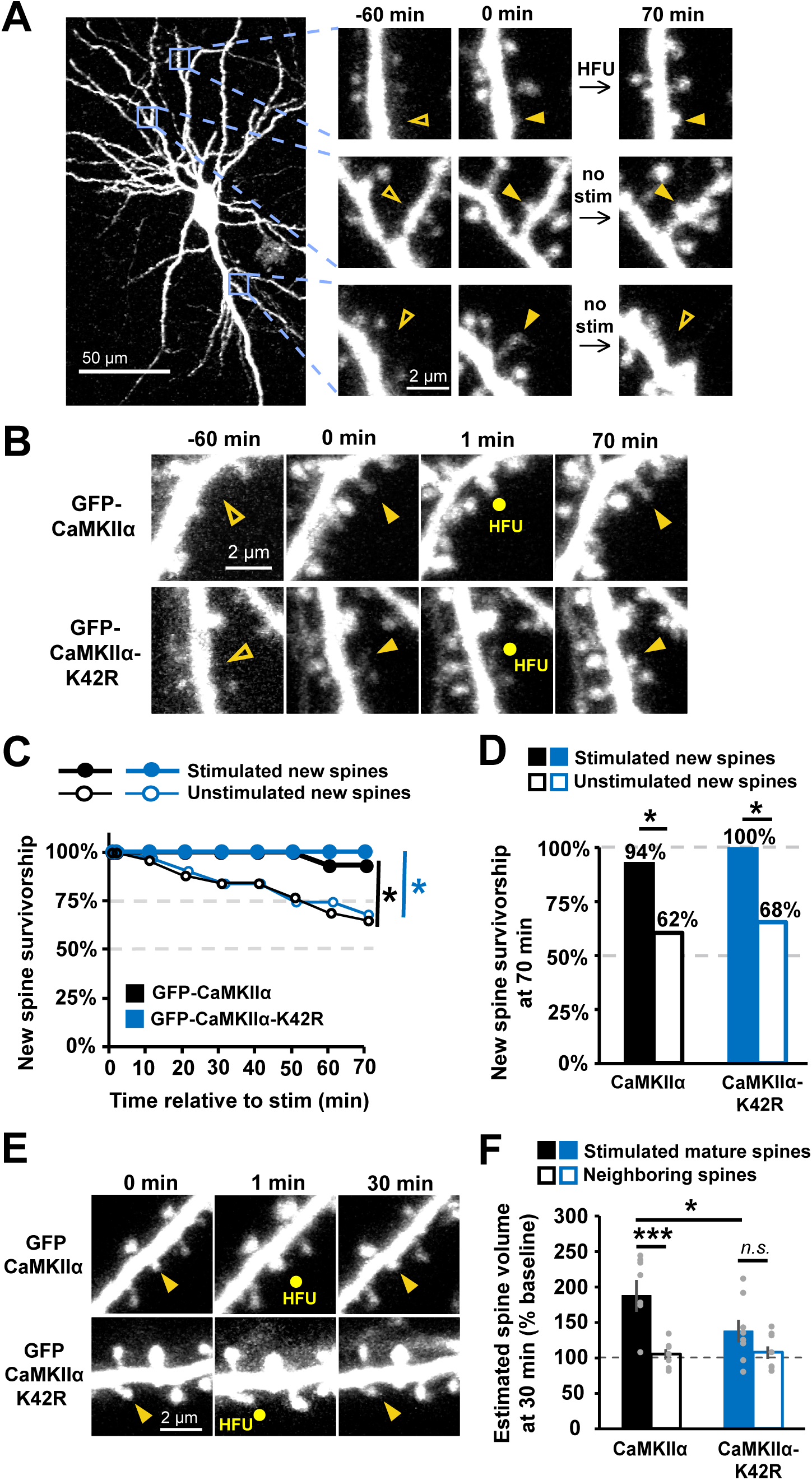
GFP-CaMKIIα-K42R does not impair activity-dependent new spine stabilization. ***(A)*** Images (red channel) of a hippocampal CA1 neuron (DIV 7-9) expressing DsRed-Express and GFP-CaMKIIα. Three new spines appeared (solid arrowheads), one of which was stimulated with HFU. One unstimulated spine was eliminated (open arrowhead). ***(B)*** Images (red channel) of dendrites from CA1 neurons expressing DsRed-Express and either GFP-CaMKIIα (top row) or GFP-CaMKIIα-K42R (bottom row) showing new dendritic spines stimulated at 1 min with HFU that were stable until 70 min. ***(C)*** HFU stimulation enhanced new spine survivorship (filled circles; WT: n=16 spines/16 cells; K42R: n=14 spines/14 cells) relative to unstimulated new spines (open circles; WT: n=31 spines/16 cells; K42R: n=32 spines/14 cells) on the same cells for both GFP-CaMKIIα-WT (black) and GFP-CaMKIIα-K42R (blue) conditions. ***(D)*** Survivorship of HFU-stimulated new spines (filled bars) at 70 min was increased compared to unstimulated new spines (open bars) on the same cells for both GFP-CaMKIIα-WT (black) and GFP-CaMKIIα-K42R (blue). ***(E)*** Images (red channel) of dendrites of CA1 neurons (DIV 7-9) before HFU at mature spines, immediately after HFU, and at 30 min after HFU. ***(F)*** GFP-CaMKIIα-K42R expression impaired HFU-induced long-term growth of mature spines (filled blue; n= 8 spines/8 cells) that is retained in cells expressing GFP-CaMKIIα (filled black; n= 8 spines/8 cells). Log-rank task in ***C***, Barnard’s exact test in ***D***, and two-way ANOVA with Bonferroni multiple comparisons test in ***F***. *p <0.05, **p < 0.01, ***p < 0.001.

While the K42R mutation acts in a dominant-negative manner, it retains residual kinase activity in response to glutamatergic stimulation (Rossetti et al., 2017; Tullis et al., 2020), and we were also concerned that transfected cells might contain fully endogenous CaMKII holoenzymes lacking the mutant subunit. Residual levels of CaMKII activity could be sufficient to promote the enzymatic interactions and signaling cascades necessary to stabilize new spines. As an independent means to test the role CaMKIIα enzymatic activity in activity-induced new spine stabilization, we pharmacologically inhibited CaMKIIα using staurosporine, a potent, broad-spectrum kinase inhibitor that competitively binds the ATP-binding pocket of CaMKIIα. Unlike many of the more widely used CaMKIIα kinase inhibitors with higher specificity, staurosporine does not interfere with the interaction between activated CaMKIIα and the GluN2B subunit (Barcomb et al., 2013). Using staurosporine to inhibit CaMKIIα thus allowed us to distinguish between the requirement for GluN2B binding (Hill and Zito, 2013) and the potential requirement for kinase activity in activity-induced new spine stabilization.

Using time-lapse 2-photon imaging of dendrites on hippocampal CA1 neurons expressing GFP, we identified multiple new spines that spontaneously grew on each cell (**Fig. 3A**). We then added staurosporine (final concentration of 1 µM) or an equivalent volume of vehicle for the remainder of the experiment. After a 30 min incubation in either staurosporine or vehicle, one new spine per cell was exposed to HFU stimulation and survivorship was monitored for stimulated and unstimulated new spines on the same cell. We found that stimulated new spines were significantly more stable than unstimulated new spines on the same cells after incubation in either vehicle (stim: 100%; unstim: 65%; *p = 0*.*04*) or staurosporine (stim: 100%; unstim: 70%; *p = 0*.*04*) (**Fig. 3B, C**). Furthermore, the rate of stimulated and unstimulated new spine survivorship was not different between the vehicle and staurosporine conditions (stim veh vs sta: *p = 0*.*99*; unstim veh vs sta: *p = 0*.*80*). Importantly, we confirmed that HFU-induced long-term growth of mature spines was blocked by staurosporine (101 ± 9%; *p = 0*.*03*) but intact in vehicle (140 ± 11%; *p = 0*.*99*), indicating the effectiveness of staurosporine as a kinase inhibitor (**Fig. 3D, E**). Our results with staurosporine are consistent with our finding that genetic inhibition of CaMKIIα kinase activity did not impair activity-induced new spine stabilization. Together, these results strongly support that CaMKIIα kinase activity is not necessary for activity-dependent new spine stabilization.

**Figure 3.**
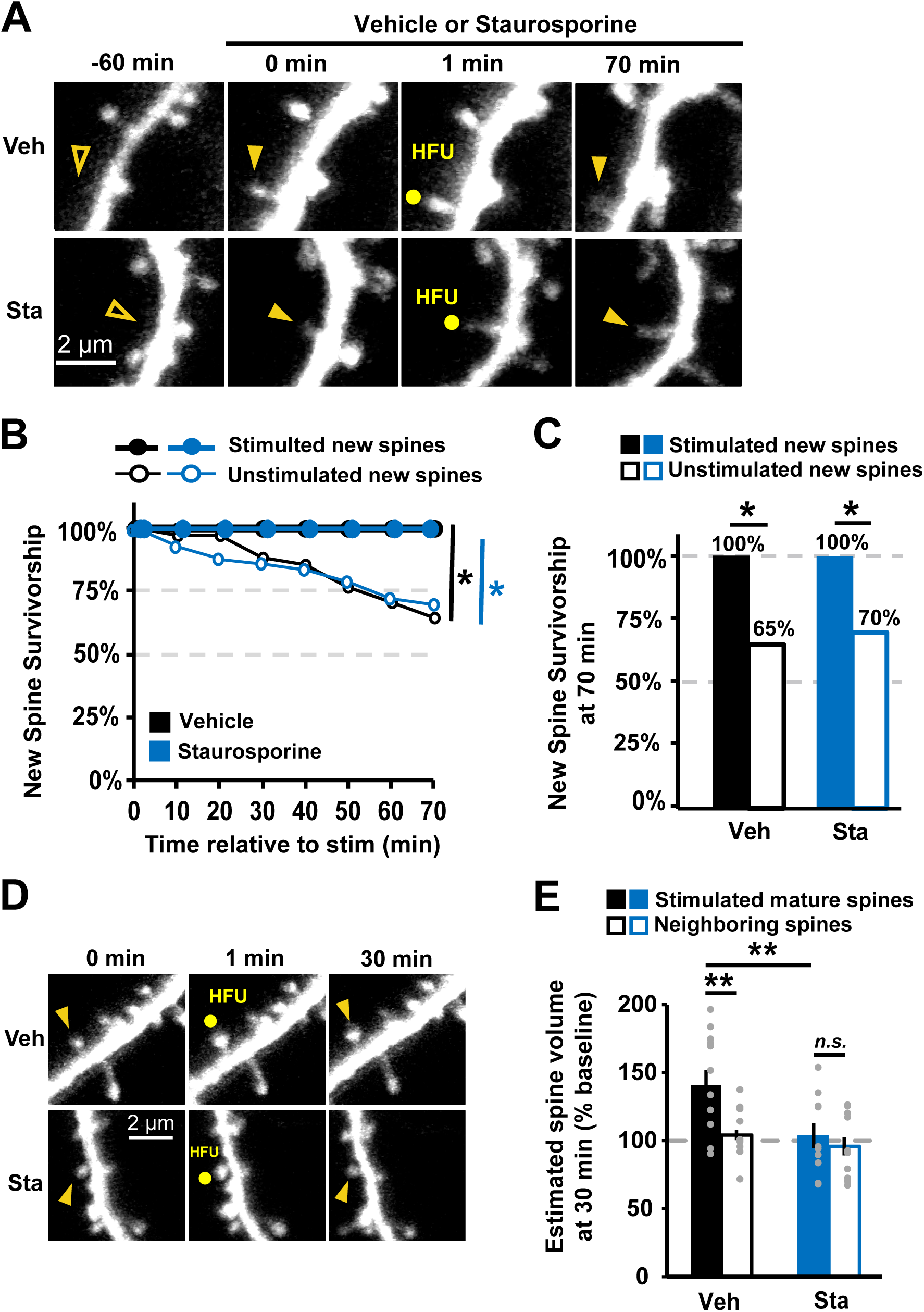
Inhibition of CaMKIIα kinase activity with staurosporine does not impair activity-dependent new spine stabilization. ***(A)*** Images (green channel) of spontaneous new spine outgrowth (filled arrowhead at 0 min) on dendrites of GFP-transfected hippocampal CA1 neurons (DIV 7-9). One new spine per neuron was stimulated with HFU following 30 min pre-incubation in either vehicle (top row) or 1 µm staurosporine (Sta; bottom row). ***(B)*** Survivorship of stimulated new spines (filled circles; veh: 9 spines/9 cells; Sta: 11 spines/11 cells) was enhanced relative to unstimulated new spines on the same cells (open circles; veh: 33 spines/9 cells; Sta: 43 spines/11 cells) in both vehicle (black) and staurosporine (blue) conditions. ***(C)*** Survivorship of HFU-stimulated new spines (filled bars) at 70 min was increased compared to unstimulated new spines (open bars) on the same cells in both vehicle (black) and staurosporine (blue) conditions. ***(D)*** Images of dendrites (green channel) on GFP-transfected CA1 neurons before HFU, immediately after HFU (yellow circle), and at 30 min. ***(E)***. Incubation with 1 µM staurosporine (sta; filled blue; n=10 spines/10 cells) blocked HFU-induced long-term growth of mature spines (101 ± 9%; *p = 0*.*03*), which was intact in vehicle conditions (veh; filled black; n=12 spines/12 cells; 140 ± 11%; *p = 0*.*99*). Volume of unstimulated neighbors was unchanged (open bars; veh: 104 ± 3%, *p = 0*.*99*; K42R: 96 ± 7%, *p = 0*.*99*). Log-rank task in ***B***, Barnard’s exact test in ***C***, and two-way ANOVA with Bonferroni multiple-comparisons test in ***E***. *p <0.05, **p < 0.01, ***p < 0.001.

### Knock-down of CaMKIIα blocks activity-dependent new spine stabilization

We next set out to test whether kinase-independent functions of CaMKIIα are required for nascent spine stabilization. Beyond its enzymatic activities, CaMKIIα plays a number of structural and scaffolding roles, independent of those performed by CaMKIIβ, most of which are facilitated by interactions with other synaptic proteins such as α-actinin, densin-180, the GluN2B subunit of the NMDAR, the proteasome, and PSD MAGUKs (Bingol et al., 2010; Krapivinsky et al., 2004; Walikonis et al., 2001). Some of these scaffolding and structural roles of CaMKIIα are distinct from its enzymatic roles and do not require CaMKIIα kinase activity (Barcomb et al., 2013; Bingol et al., 2010; Krapivinsky et al., 2004; Pi et al., 2010). These interactions would require precise regulation of the amounts of available CaMKIIα and its physical interactions with potential binding partners. Thus, decreased levels of endogenous CaMKIIα would likely interfere with these structural and scaffolding activities, some of which may be necessary for activity-dependent new spine stabilization.

We tested whether structural and/or scaffolding activities of CaMKIIα are needed to support activity-dependent nascent spine stabilization using an shRNA-mediated knock-down of endogenous CaMKIIα with an shRNA that was designed and validated in previous work (Lemieux et al., 2012). We validated this CaMKIIα-shRNA in our preparation by demonstrating that HFU-induced long-term growth of mature spines was blocked by knock-down of CaMKIIα (98 ± 10%; *p = 0*.*99*) and rescued by co-expression of shRNA-resistant GFP-CaMKIIα* (200 ± 26%; *p = 0*.*04*; **Fig. 4A, B**). We next applied this CaMKIIα-shRNA to test HFU-induced new spine stabilization. We found that knock-down of CaMKIIα disrupted HFU-induced new spine stabilization (**Fig. 4C-E**; stim: 60%, unstim: 61%; *p = 0*.*98*). Interestingly, new spines on cells expressing CaMKIIα-shRNA appeared to be initially stabilized by HFU stimulation, but only transiently, as new spine survivorship at 70 min was the same as for unstimulated spines (**Fig. 4D**). To rule out possible effects of non-specific shRNA activity, we rescued the knock-down by co-expressing an shRNA-resistant form of GFP-CaMKIIα (GFP-CaMKIIα*). Rescuing CaMKIIα levels restored activity-dependent new spine stabilization, as new spines that received the HFU stimulus were again significantly more stable than unstimulated control new spines (**Fig. 4C-E**; stim: 100%, unstim: 67%; *p = 0*.*03*). These results confirm a role CaMKIIα in activity-induced new spine stabilization. Together with our previous results, we conclude that non-enzymatic CaMKIIα function is required to enhance new spine stabilization downstream of strong glutamatergic stimulation.

**Figure 4.**
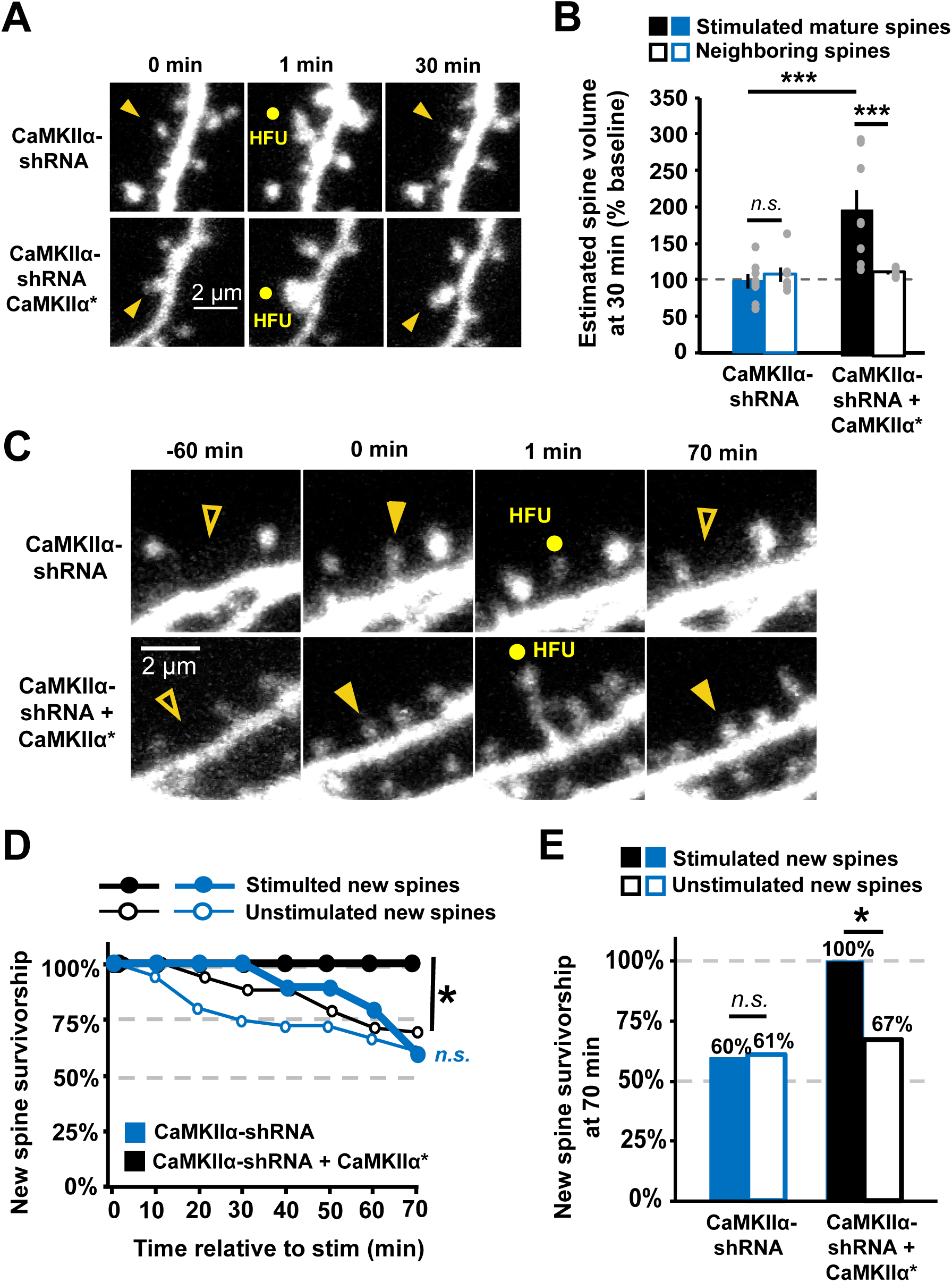
CaMKIIα knock-down blocks activity-dependent new spine stabilization. ***(A)*** Images of dendrites on organotypic hippocampal CA1 neurons (DIV 7-9) before HFU, immediately after HFU (yellow circle), and at the final 30 min time point. ***(B)*** shRNA-mediated knockdown of CaMKIIα (filled blue; n=8 spines/8 cells) impaired HFU-induced long-term spine growth that was restored with co-expression of shRNA-resistant GFP-CaMKIIα* (black; n=8 spines/8 cells). ***(C)*** Images of dendrites (red channel) on CA1 neurons (DIV 7-9) expressing a DsRed-Express cell fill and either CaMKIIα shRNA (top) or CaMKIIα shRNA + shRNA-resistant GFP-CaMKIIα* (bottom). Spontaneous new spine outgrowth (filled arrowheads) was observed in both conditions. HFU-induced new spine stabilization failed following knock-down of CaMKIIα (open arrowhead at 70 min), but was rescued with expression of shRNA-resistant GFP-CaMKIIα*. ***(D)*** Knockdown of CaMKIIα disrupted the stabilization of new spines (filled circles) as compared with unstimulated new spines (open circles) at times beyond 30-40 min (blue; stimulated: 10 spines/10 cells; unstimulated: 36 spines, 10 cells). Rescuing CaMKIIα restored activity-dependent new spine stabilization (black; stim: 11 spines/11 cells; unstim: 53 spines/11 cells). ***(E)*** Activity-dependent new spine stabilization at 70 min (filled bars) was not significantly different from that of unstimulated new spines (open bars) following knockdown of CaMKIIα (black) but was restored when CaMKIIα is rescued with shRNA-resistant CaMKIIα* (blue). Two-way ANOVA with Bonferroni multiple-comparisons test in ***B***, log-rank task in ***D***, and Barnard’s exact test in ***E***. *p <0.05, **p < 0.01, ***p < 0.001.

### Overexpression of pseudo-autophosphorylated CaMKIIα enhances basal spine survivorship independent of kinase activity

We next probed whether CaMKIIα’s non-enzymatic structural and/or scaffolding activities are not only necessary, but sufficient to enhance activity-dependent new spine stabilization. We took advantage of phospho-mimetic CaMKIIα mutants that increase basal levels of CaMKIIα-GluN2B binding (Barcomb et al., 2014), specifically the replacement of threonine 286 with an aspartic acid, or CaMKIIα-T286D (Pi et al., 2010). The T286D mutation renders CaMKIIα constitutively active, allowing interactors and substrates access to the kinase and regulatory domains. Pairing this mutation with the K42R point mutation generates increased autonomous CaMKIIα interactions, while blocking CaMKIIα enzymatic activities.

To determine the effect of autonomous CaMKIIα on new spine survivorship with and without kinase activity, we expressed the GFP-CaMKIIα-T286D or GFP-CaMKIIα-K42R/T286D with a dsRed-Express cell fill in organotypic hippocampal slice cultures. For these experiments where we wanted to determine whether constitutively autonomous CaMKIIα was sufficient to enhance new spine survivorship, we did not expose new spines to our HFU protocol, instead we monitored basal new spine stability over a period of 70 min using time-lapse imaging. We found that new spines were more stable on cells expressing either GFP-CaMKIIα-T286D or GFP-CaMKIIα-K42R/T286D compared to new spines on cells expressing only DsRed-Express (**Fig 5**; DsRed: 63%; T286D: 83%, *p = 0*.*02*; K42R/T286D: 85%, *p = 0*.*01*). Importantly, expression of wild-type (WT) GFP-CaMKIIα did not alter new spine survivorship as compared to dsRed-

**Figure 5.**
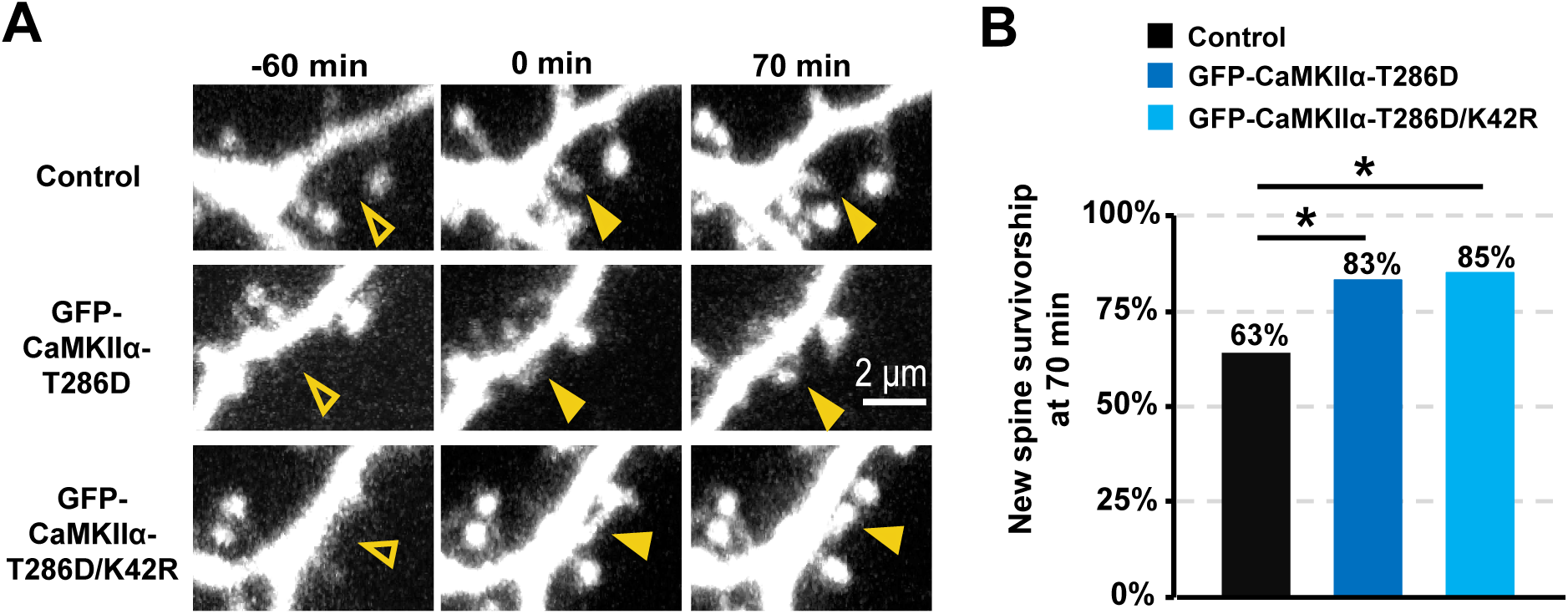
Overexpression of constitutively autonomous CaMKIIα enhances basal spine survivorship independent of kinase activity. ***(A)*** Images (red channel) showing spontaneous new spine outgrowth (filled arrowhead at 0 min) and stabilization (filled arrowhead at 70 min) on dendrites of hippocampal neurons (DIV 7-9) expressing DsRed-Express alone (top row) or co-expressed with GFP-CaMKIIα-T286D (middle row) or GFP-CaMKIIα-K42R/T286D (bottom row). ***(B)*** Basal spine survivorship rates were higher on cells expressing GFP-CaMKIIα-T286D (dark blue; 63 spines/ 10 cells) and GFP-CaMKIIα-K42R/T286D (light blue; 53 spines/ 7 cells) compared to survivorship rates on cells expressing only DsRed-Express (black; 51 spines/ 10 cells). Barnard’s exact test with Bonferroni multiple-comparisons correction. *p <0.05, **p < 0.01, ***p < 0.001.

Express alone (DsRed: 67%; WT: 65%; *p = 0*.*84*), so increased survivorship was due to pseudo-autophosphorylated CaMKIIα, independent of kinase activity. Our findings support a model in which CaMKIIα-GluN2B binding facilitates non-enzymatic CaMKIIα functions that are both necessary and sufficient for enhancing new spine stabilization.

## DISCUSSION

### Molecular composition of nascent dendritic spines

There is substantial evidence indicating that the formation of new spines and their ability to persist and integrate into functional synaptic circuits is crucial to learning (Albarran et al., 2021; Hayashi-Takagi et al., 2015; Roberts et al., 2010; Xu et al., 2009; Yang et al., 2009). Despite this vital role, the molecular composition and signaling pathways at play in new spines remain largely unexplored. New spines do share some molecular and functional properties with mature spines; new spine AMPAR currents are comparable to those recorded from mature spines of similar size (Kwon and Sabatini, 2011; Zito et al., 2009) and ultrastructural evidence shows that a subset of new spines are found directly apposed to presynaptic boutons (Knott et al., 2006; Trachtenberg et al., 2002; Zito et al., 2009), suggesting that new spines are rapidly equipped to respond to glutamatergic stimulation and incorporated into neural circuits.

Still, new spines differ from mature spines in several key ways. Most notably, the very low expression levels in new spines of the PSD-family MAGUKs (De Roo et al., 2008a; Lambert et al., 2017), key scaffolding molecules that regulate synaptic strength, maturation, and stability (Boehm et al., 2006; Cane et al., 2014; Ehrlich and Malinow, 2004; Elias et al., 2008; Taft and Turrigiano, 2014). NMDAR currents are also smaller in new spines (Kwon and Sabatini, 2011; Zito et al., 2009), where they demonstrate greater diffusional coupling to the dendrite (Zito et al., 2009). PSD-family MAGUKS, NMDAR-mediated signaling, and spine morphologies associated with a high degree of compartmentalization are all thought to regulate synaptic stability (Cane et al., 2014; De Roo et al., 2008a; De Roo et al., 2008b; Lambert et al., 2017; Taft and Turrigiano, 2014), suggesting that the low basal survivorship rates of new spines may be due to their distinct molecular composition and signaling. Identifying the molecular signaling pathways at play in new spines is therefore crucial to understand the mechanisms involved in their stabilization.

Here, we show that, unlike GFP-tagged PSD-family MAGUKs, new spines express GFP-CaMKIIα at levels comparable to those in mature spine levels, independent of CaMKIIα kinase activity. CaMKIIα’s presence in new spines supports that CaMKIIα signaling could play a critical role in new spine function. Indeed, evidence supports a requirement for the CaMKIIα-GluN2B interaction not only in activity-dependent new spine stabilization (Hill and Zito, 2013), but also in spontaneous and activity-dependent new spine outgrowth (Hamilton et al., 2012), suggesting that CaMKIIα’s functions at new spines may precede any form of synaptic stimulation.

### Role of CaMKIIα kinase activity in new spine stabilization

Despite our finding that CaMKIIα is present at mature levels in new spines, we were surprised to find that CaMKIIα kinase activity is not required for enhanced new spine stabilization induced either by strong glutamatergic stimulation at single spines or by overexpression of the CaMKIIα-K42R/T286 phospho-mutant. Our results in new spines are in contrast with what is known regarding the important role of CaMKIIα kinase activity in stabilization of the long-term growth of mature spines (Araki et al., 2015; Cornelia Koeberle et al., 2017). However, major changes to the molecular composition of new spines occur during the maturation process, including the recruitment of PSD-family MAGUKs (De Roo et al., 2008a; Lambert et al., 2017), no doubt creating a vastly different biochemical signaling environment in the new spine as it develops.

Indeed, it is possible that, while CaMKIIα kinase activity is not required to enhance new spine stabilization on the time scale of 70-130 min after new spine growth, as we observed in our experiments, it may be necessary at later times, for example following the delayed recruitment of other synaptic proteins, such as PSD-family MAGUKS. Notably, on the very earliest time scales (<40-100 min), even non-enzymatic CaMKIIα function does not appear to be required for activity-dependent new spine stabilization, as we show that stimulated new spines are initially stabilized even on cells where CaMKIIα is knocked down. Overall, our data demonstrate that CaMKIIα kinase function is not required for the early steps of new spine stabilization, within the first few hours following new spine outgrowth.

### Role of GluN2B-CaMKIIα binding in new spine stabilization

Our finding that CaMKIIα kinase activity is not required for activity-dependent new spine stabilization leaves an undefined role for the required CaMKIIα-GluN2B interaction. This interaction has long been known to be important in the regulation of basal synaptic transmission and LTP maintenance (Barcomb et al., 2014; Barria and Malinow, 2002; Halt et al., 2012; Incontro et al., 2018), where it is thought to play a role in bringing Ca^2+^/CaM-activated CaMKII closer to its targeted substrates to alter synaptic transmission and synaptic strengthening in a kinase-dependent manner. In new spines, our results instead support a non-enzymatic role for CaMKIIα in new spine stabilization. Indeed, we show that, although CaMKIIα kinase activity is not required, knockdown of CaMKIIα disrupts activity-dependent new spine stabilization. Altogether our results suggest that the interaction between GluN2B and CaMKIIα is required to support a primarily structural or scaffolding role for CaMKIIα in new spine stabilization.

While we found that CaMKIIα-T286D, which enhanced the interaction between GluN2B and CaMKIIα, also increased basal spine stabilization, survivorship rates for CaMKIIα-T286D were lower than observed for new spines that received HFU stimulation. At mature spines glutamatergic stimulation initiates a number of concurrent signaling cascades and molecular changes, such as NMDAR and mGluR activation and downstream signaling mechanisms (Bosch et al., 2014; Lee et al., 2003; Malinow, 2003; Murakoshi et al., 2011; Stein et al., 2021) that are not replicated by overexpression of the CaMKIIα-T286D phospho-mutant. It is likely that at least a subset of these mechanisms acts in conjunction with GluN2B-CaMKIIα binding to enhance activity-dependent new spine stabilization. In addition, the GluN2B-CaMKIIα interaction may serve to bring CaMKIIβ, which complexes with CaMKIIα at a 3:9 ratio in the hippocampus, within optimal proximity to its binding partners in order to regulate cytoskeletal stability (Kim et al., 2015; Kim et al., 2019; Okamoto et al., 2007).

### Non-enzymatic CaMKIIα function in new spine stabilization

Although CaMKIIβ is perhaps more well recognized than CaMKIIα for its non-enzymatic functions in regulating the spine actin cytoskeleton, CaMKIIα has also been shown to participate in several functionally important scaffolding and structural interactions that are distinct from those made by CaMKIIβ (Bayer and Schulman, 2019; Hell, 2014). Some of these interactions are less likely to be relevant for the earliest stages of new spine stabilization, such as roles with stargazin, TARP γ-8, or the Rac-1 activating RAKEC (Opazo et al., 2010; Opazo et al., 2012; Park et al., 2016; Saneyoshi et al., 2019), as they require either binding to PSD-family MAGUKs or CaMKIIα kinase activity. However, other known CaMKIIα interactions are independent of these requirements and therefore would make attractive candidates for roles in new spine stabilization, such as the activity-dependent binding of CaMKIIα directly to the 26S proteasome, and indirect interactions of CaMKIIα with SynGAP-1α via the multi-PDZ domain protein MUPP-1 (Bingol et al., 2010; Krapivinsky et al., 2004).

Indeed, CaMKIIα’s non-enzymatic interactions with the proteasome and the MUPP1-SynGAP-1α complex appear particularly promising in the context of understanding new spine stabilization. SynGAP-1α is a negative-regulator of synapse maturation, and its exclusion from synapses contributes to synaptic strengthening, precocious PSD-95 accumulation, and increased spine volume (Aceti et al., 2015; Araki et al., 2015; Clement et al., 2012; Vazquez et al., 2004). Interestingly, activity-dependent dissociation of the MUPP1-SynGAP-1α complex from CaMKIIα does not require CaMKIIα kinase activity (Krapivinsky et al., 2004) and thus may provide a mechanism for SynGAP-1α dispersion (Araki et al., 2015) from new spines, independent of PSD-family MAGUKs and kinase activity. Furthermore, activity-dependent new spine formation requires the proteasome (Hamilton et al., 2012), which may remain accumulated at sites of new spine formation, where it could play a role in the activity-dependent degradation of negative regulators of synapse stability and maturation. Indeed, there is evidence that the proteasome mediates degradation of SynGAP (Zhang et al., 2020) and Ephexin5 (Hamilton et al., 2017), which both have roles in regulating dendritic spine stability.

The elucidation of the role of these two proteins and of other non-enzymatic functions of CaMKIIα downstream of GluN2B-CaMKIIα to promote new spine survivorship is an intriguing and compelling avenue for future study.

## Acknowledgements

This work was supported by the National Institutes of Health (R01 NS062736, T32 MH112507) and a grant from the Howard Hughes Medical Institute through the Gilliam Fellowships for Advanced Study (N.C.). We thank Paul De Koninck for kindly sharing the mEGFP-tagged CaMKIIα-WT, -K42R, -T286D, and -K42R/T286D, CaMKIIα-shRNA, and shRNA-resistant CaMKIIα* constructs. We thank M.Anisimova, S.Petshow, J.Flores, M. Ferns, J.Gray, J.Hell, and K.McAllister for critical input and/or reading of the manuscript.

## AUTHOR CONTRIBUTIONS

N.C. and K.Z. conceived the study and designed the experiments. N.C. performed the experiments, analyzed the data, and wrote the first draft of the manuscript. N.C. and K.Z. edited the manuscript.

## REFERENCES

Aceti, M., Creson, T.K., Vaissiere, T., Rojas, C., Huang, W.C., Wang, Y.X., Petralia, R.S., Page, D.T., Miller, C.A., and Rumbaugh, G. (2015). Syngap1 haploinsufficiency damages a postnatal critical period of pyramidal cell structural maturation linked to cortical circuit assembly. Biol Psychiatry 77, 805–815.

Albarran, E., Raissi, A., Jaidar, O., Shatz, C.J., and Ding, J.B. (2021). Enhancing motor learning by increasing the stability of newly formed dendritic spines in the motor cortex. Neuron 109, 3298–3311 e3294.

Araki, Y., Zeng, M., Zhang, M., and Huganir, R.L. (2015). Rapid dispersion of SynGAP from synaptic spines triggers AMPA receptor insertion and spine enlargement during LTP. Neuron 85, 173–189.

Barcomb, K., Buard, I., Coultrap, S.J., Kulbe, J.R., O’Leary, H., Benke, T.A., and Bayer, K.U. (2014). Autonomous CaMKII requires further stimulation by Ca2+/calmodulin for enhancing synaptic strength. FASEB J 28, 3810–3819.

Barcomb, K., Coultrap, S.J., and Bayer, K.U. (2013). Enzymatic activity of CaMKII is not required for its interaction with the glutamate receptor subunit GluN2B. Mol Pharmacol 84, 834–843.

Barria, A., and Malinow, R. (2002). Subunit-specific NMDA receptor trafficking to synapses. Neuron 35, 345–353.

Bayer, K.U., and Schulman, H. (2019). CaM Kinase: Still Inspiring at 40. Neuron 103, 380–394.

Berry, K.P., and Nedivi, E. (2017). Spine Dynamics: Are They All the Same? Neuron 96, 43–55.

Bingol, B., Wang, C.F., Arnott, D., Cheng, D., Peng, J., and Sheng, M. (2010). Autophosphorylated CaMKIIalpha acts as a scaffold to recruit proteasomes to dendritic spines. Cell 140, 567–578.

Boehm, J., Ehrlich, I., Hsieh, H., and Malinow, R. (2006). Two mutations preventing PDZ-protein interactions of GluR1 have opposite effects on synaptic plasticity. Learn Mem 13, 562–565.

Bosch, M., Castro, J., Saneyoshi, T., Matsuno, H., Sur, M., and Hayashi, Y. (2014). Structural and molecular remodeling of dendritic spine substructures during long-term potentiation. Neuron 82, 444–459.

Cane, M., Maco, B., Knott, G., and Holtmaat, A. (2014). The relationship between PSD-95 clustering and spine stability in vivo. J Neurosci 34, 2075–2086.

Clement, J.P., Aceti, M., Creson, T.K., Ozkan, E.D., Shi, Y., Reish, N.J., Almonte, A.G., Miller, B.H., Wiltgen, B.J., Miller, C.A., et al. (2012). Pathogenic SYNGAP1 mutations impair cognitive development by disrupting maturation of dendritic spine synapses. Cell 151, 709–723.

Cornelia Koeberle, S., Tanaka, S., Kuriu, T., Iwasaki, H., Koeberle, A., Schulz, A., Helbing, D.L., Yamagata, Y., Morrison, H., and Okabe, S. (2017). Developmental stage-dependent regulation of spine formation by calcium-calmodulin-dependent protein kinase IIalpha and Rap1. Sci Rep 7, 13409.

De Roo, M., Klauser, P., Mendez, P., Poglia, L., and Muller, D. (2008a). Activity-dependent PSD formation and stabilization of newly formed spines in hippocampal slice cultures. Cereb Cortex 18, 151–161.

De Roo, M., Klauser, P., and Muller, D. (2008b). LTP promotes a selective long-term stabilization and clustering of dendritic spines. PLoS Biol 6, e219.

Ehrlich, I., and Malinow, R. (2004). Postsynaptic density 95 controls AMPA receptor incorporation during long-term potentiation and experience-driven synaptic plasticity. J Neurosci 24, 916–927.

Elias, G.M., Elias, L.A., Apostolides, P.F., Kriegstein, A.R., and Nicoll, R.A. (2008). Differential trafficking of AMPA and NMDA receptors by SAP102 and PSD-95 underlies synapse development. Proc Natl Acad Sci U S A 105, 20953–20958.

Halt, A.R., Dallapiazza, R.F., Zhou, Y., Stein, I.S., Qian, H., Juntti, S., Wojcik, S., Brose, N., Silva, A.J., and Hell, J.W. (2012). CaMKII binding to GluN2B is critical during memory consolidation. EMBO J 31, 1203–1216.

Hamilton, A.M., Lambert, J.T., Parajuli, L.K., Vivas, O., Park, D.K., Stein, I.S., Jahncke, J.N., Greenberg, M.E., Margolis, S.S., and Zito, K. (2017). A dual role for the RhoGEF Ephexin5 in regulation of dendritic spine outgrowth. Mol Cell Neurosci 80, 66–74.

Hamilton, A.M., Oh, W.C., Vega-Ramirez, H., Stein, I.S., Hell, J.W., Patrick, G.N., and Zito, K. (2012). Activity-dependent growth of new dendritic spines is regulated by the proteasome. Neuron 74, 1023–1030.

Hayashi-Takagi, A., Yagishita, S., Nakamura, M., Shirai, F., Wu, Y.I., Loshbaugh, A.L., Kuhlman, B., Hahn, K.M., and Kasai, H. (2015). Labelling and optical erasure of synaptic memory traces in the motor cortex. Nature 525, 333–338.

Hell, J.W. (2014). CaMKII: claiming center stage in postsynaptic function and organization. Neuron 81, 249–265.

Hill, T.C., and Zito, K. (2013). LTP-induced long-term stabilization of individual nascent dendritic spines. J Neurosci 33, 678–686.

Holtmaat, A.J., Trachtenberg, J.T., Wilbrecht, L., Shepherd, G.M., Zhang, X., Knott, G.W., and Svoboda, K. (2005). Transient and persistent dendritic spines in the neocortex in vivo. Neuron 45, 279–291.

Incontro, S., Diaz-Alonso, J., Iafrati, J., Vieira, M., Asensio, C.S., Sohal, V.S., Roche, K.W., Bender, K.J., and Nicoll, R.A. (2018). The CaMKII/NMDA receptor complex controls hippocampal synaptic transmission by kinase-dependent and independent mechanisms. Nat Commun 9, 2069.

Kasai, H., Ziv, N.E., Okazaki, H., Yagishita, S., and Toyoizumi, T. (2021). Spine dynamics in the brain, mental disorders and artificial neural networks. Nat Rev Neurosci 22, 407–422.

Kim, K., Lakhanpal, G., Lu, H.E., Khan, M., Suzuki, A., Hayashi, M.K., Narayanan, R., Luyben, T.T., Matsuda, T., Nagai, T., et al. (2015). A Temporary Gating of Actin Remodeling during Synaptic Plasticity Consists of the Interplay between the Kinase and Structural Functions of CaMKII. Neuron 87, 813–826.

Kim, K., Suzuki, A., Kojima, H., Kawamura, M., Miya, K., Abe, M., Yamada, I., Furuse, T., Wakana, S., Sakimura, K., et al. (2019). Autophosphorylation of F-actin binding domain of CaMKIIbeta is required for fear learning. Neurobiol Learn Mem 157, 86–95.

Knott, G.W., Holtmaat, A., Wilbrecht, L., Welker, E., and Svoboda, K. (2006). Spine growth precedes synapse formation in the adult neocortex in vivo. Nat Neurosci 9, 1117–1124.

Krapivinsky, G., Medina, I., Krapivinsky, L., Gapon, S., and Clapham, D.E. (2004). SynGAP-MUPP1-CaMKII synaptic complexes regulate p38 MAP kinase activity and NMDA receptor-dependent synaptic AMPA receptor potentiation. Neuron 43, 563–574.

Kwon, H.B., and Sabatini, B.L. (2011). Glutamate induces de novo growth of functional spines in developing cortex. Nature 474, 100–104.

Lambert, J.T., Hill, T.C., Park, D.K., Culp, J.H., and Zito, K. (2017). Protracted and asynchronous accumulation of PSD95-family MAGUKs during maturation of nascent dendritic spines. Dev Neurobiol 77, 1161–1174.

Lee, H.K., Takamiya, K., Han, J.S., Man, H., Kim, C.H., Rumbaugh, G., Yu, S., Ding, L., He, C., Petralia, R.S., et al. (2003). Phosphorylation of the AMPA receptor GluR1 subunit is required for synaptic plasticity and retention of spatial memory. Cell 112, 631–643.

Lemieux, M., Labrecque, S., Tardif, C., Labrie-Dion, E., Lebel, E., and De Koninck, P. (2012). Translocation of CaMKII to dendritic microtubules supports the plasticity of local synapses. J Cell Biol 198, 1055–1073.

Malinow, R. (2003). AMPA receptor trafficking and long-term potentiation. Philos Trans R Soc Lond B Biol Sci 358, 707–714.

Matsuzaki, M., Honkura, N., Ellis-Davies, G.C., and Kasai, H. (2004). Structural basis of long-term potentiation in single dendritic spines. Nature 429, 761–766.

Murakoshi, H., Wang, H., and Yasuda, R. (2011). Local, persistent activation of Rho GTPases during plasticity of single dendritic spines. Nature 472, 100–104.

Okamoto, K., Narayanan, R., Lee, S.H., Murata, K., and Hayashi, Y. (2007). The role of CaMKII as an F-actin-bundling protein crucial for maintenance of dendritic spine structure. Proc Natl Acad Sci U S A 104, 6418–6423.

Opazo, P., Labrecque, S., Tigaret, C.M., Frouin, A., Wiseman, P.W., De Koninck, P., and Choquet, D. (2010). CaMKII triggers the diffusional trapping of surface AMPARs through phosphorylation of stargazin. Neuron 67, 239–252.

Opazo, P., Sainlos, M., and Choquet, D. (2012). Regulation of AMPA receptor surface diffusion by PSD-95 slots. Curr Opin Neurobiol 22, 453–460.

Opitz-Araya, X., and Barria, A. (2011). Organotypic hippocampal slice cultures. J Vis Exp.

Park, J., Chavez, A.E., Mineur, Y.S., Morimoto-Tomita, M., Lutzu, S., Kim, K.S., Picciotto, M.R., Castillo, P.E., and Tomita, S. (2016). CaMKII Phosphorylation of TARPgamma-8 Is a Mediator of LTP and Learning and Memory. Neuron 92, 75–83.

Pi, H.J., Otmakhov, N., El Gaamouch, F., Lemelin, D., De Koninck, P., and Lisman, J. (2010). CaMKII control of spine size and synaptic strength: role of phosphorylation states and nonenzymatic action. Proc Natl Acad Sci U S A 107, 14437–14442.

Pologruto, T.A., Sabatini, B.L., and Svoboda, K. (2003). ScanImage: flexible software for operating laser scanning microscopes. Biomed Eng Online 2, 13.

Roberts, T.F., Tschida, K.A., Klein, M.E., and Mooney, R. (2010). Rapid spine stabilization and synaptic enhancement at the onset of behavioural learning. Nature 463, 948–952.

Rossetti, T., Banerjee, S., Kim, C., Leubner, M., Lamar, C., Gupta, P., Lee, B., Neve, R., and Lisman, J. (2017). Memory Erasure Experiments Indicate a Critical Role of CaMKII in Memory Storage. Neuron 96, 207–216 e202.

Saneyoshi, T., Matsuno, H., Suzuki, A., Murakoshi, H., Hedrick, N.G., Agnello, E., O’Connell, R., Stratton, M.M., Yasuda, R., and Hayashi, Y. (2019). Reciprocal Activation within a Kinase-Effector Complex Underlying Persistence of Structural LTP. Neuron 102, 1199–1210 e1196.

Sanhueza, M., and Lisman, J. (2013). The CaMKII/NMDAR complex as a molecular memory. Mol Brain 6, 10.

Stein, I.S., Park, D.K., Claiborne, N., and Zito, K. (2021). Non-ionotropic NMDA receptor signaling gates bidirectional structural plasticity of dendritic spines. Cell Rep 34, 108664.

Stoppini, L., Buchs, P.A., and Muller, D. (1991). A simple method for organotypic cultures of nervous tissue. J Neurosci Methods 37, 173–182.

Taft, C.E., and Turrigiano, G.G. (2014). PSD-95 promotes the stabilization of young synaptic contacts. Philos Trans R Soc Lond B Biol Sci 369, 20130134.

Trachtenberg, J.T., Chen, B.E., Knott, G.W., Feng, G., Sanes, J.R., Welker, E., and Svoboda, K. (2002). Long-term in vivo imaging of experience-dependent synaptic plasticity in adult cortex. Nature 420, 788–794.

Tullis, J.E., Rumian, N.L., Brown, C.N., and Bayer, K.U. (2020). The CaMKII K42M and K42R mutations are equivalent in suppressing kinase activity and targeting. PLoS One 15, e0236478.

Vazquez, L.E., Chen, H.J., Sokolova, I., Knuesel, I., and Kennedy, M.B. (2004). SynGAP regulates spine formation. J Neurosci 24, 8862–8872.

Walikonis, R.S., Oguni, A., Khorosheva, E.M., Jeng, C.J., Asuncion, F.J., and Kennedy, M.B. (2001). Densin-180 forms a ternary complex with the (alpha)-subunit of Ca2+/calmodulin-dependent protein kinase II and (alpha)-actinin. J Neurosci 21, 423–433.

Woods, G., and Zito, K. (2008). Preparation of gene gun bullets and biolistic transfection of neurons in slice culture. J Vis Exp.

Woods, G.F., Oh, W.C., Boudewyn, L.C., Mikula, S.K., and Zito, K. (2011). Loss of PSD-95 enrichment is not a prerequisite for spine retraction. J Neurosci 31, 12129–12138.

Xu, T., Yu, X., Perlik, A.J., Tobin, W.F., Zweig, J.A., Tennant, K., Jones, T., and Zuo, Y. (2009). Rapid formation and selective stabilization of synapses for enduring motor memories. Nature 462, 915–919.

Yamagata, Y., Kobayashi, S., Umeda, T., Inoue, A., Sakagami, H., Fukaya, M., Watanabe, M., Hatanaka, N., Totsuka, M., Yagi, T., et al. (2009). Kinase-dead knock-in mouse reveals an essential role of kinase activity of Ca2+/calmodulin-dependent protein kinase IIalpha in dendritic spine enlargement, long-term potentiation, and learning. J Neurosci 29, 7607–7618.

Yang, G., Pan, F., and Gan, W.B. (2009). Stably maintained dendritic spines are associated with lifelong memories. Nature 462, 920–924.

Zhang, Q., Yang, H., Gao, H., Liu, X., Li, Q., Rong, R., Liu, Z., Wei, X.E., Kong, L., Xu, Y., et al. (2020). PSD-93 Interacts with SynGAP and Promotes SynGAP Ubiquitination and Ischemic Brain Injury in Mice. Transl Stroke Res 11, 1137–1147.

Zito, K., Scheuss, V., Knott, G., Hill, T., and Svoboda, K. (2009). Rapid functional maturation of nascent dendritic spines. Neuron 61, 247–258.

